# Evaluation of Micronuclei Frequency in Both Shelter and Family Cats and Dogs

**DOI:** 10.1101/2020.07.15.204230

**Authors:** Alfredo Santovito, Martina Buglisi, Manuel Scarfo’

**Author notes:** Corresponding Author: Alfredo SANTOVITO, University of Turin, Department of Life Sciences and Systems Biology, Via Accademia Albertina n. 13, 10123 – Torino (Italy).

## Abstract

Each year a lot of animals are cared for in shelters in Italy. Many of these animals have received minimal or no prior healthcare. Thus, the beneficial role animal shelters play is undeniable. Nonetheless, even well-run shelters lack the necessary resources to provide adequate conditions. It is common knowledge that group-housing can increase stress levels in family cats (*Felis silvestris lybica*) and dogs (*Canis lupus familiaris*) contributing to the development of infectious diseases and/or behavioural problems.

The aim of this study is to evaluate, through the buccal micronucleus assay, the level of genomic damage in shelter cats and dogs and compare it with that of family cats and dogs. The hypothesis is that environmental conditions such as those potentially present in shelters can affect the level of genomic damage.

The study population included thirty healthy mixed breed cats and dogs, randomly sampled, with at a minimum two-year presence in a shelter. The control group consisted of thirty healthy cats and dogs living in a home environment, using age/sex matching. The micronucleus assay was performed on one thousand exfoliated buccal mucosa cells per subject and standardized protocols were used for stress score tests.

Significant differences were found between shelter and family cats and dogs in terms of micronuclei, indicating that a condition of stress found in sheltered cats may increase the levels of genomic damage. Conversely, no significant differences in the frequency of micronuclei were found between the sexes, as well as no correlation was found between age and the frequencies of the used genomic markers.

## 1. Introduction

It is common knowledge that many animal shelters can be potentially stressful places for animals, mainly due to space restrictions, lack of resources and high animal turnover which, with overcrowding, leads to increased transmission of different pathogens (Kessler & Turner, 1999; Wells et al., 2002; Cohn, 2011). Moreover, euthanasia on cats and dogs in shelters is forbidden in Italy. As a consequence, this “no-kill policy” extends their stay in shelters, increasing the number of animals housed (Righi et al., 2019).

Undoubtedly, arriving at a shelter can be extremely stressful and even traumatic for an animal. Losing an emotional bond, changing daily routines and being placed in a different environment full of new and unusual stimuli are all conditions that often result in minimal possibilities of interaction with conspecifics and humans (Hennessy et al., 2001; Coppola et al., 2006). The lack of social interaction, the limited possibility of movement, the minimal control over the surrounding environment and the unpredictable noise levels can make living in a shelter a stressful condition, particularly, for extremely social animals such as dogs (Beerda et al., 2000; Wells et al., 2002; Taylor et al., 2007; Titulaer et al., 2013). For example, it was observed that staying in a shelter can induce behavioural changes in dogs as well as significantly modify their behaviour (Wells & Hepper, 2000; Cozzi et al., 2016). An increased frequency of autogrooming, circling, eating faeces, paw lifting, standing upright, digging, whining, and scratching are all examples of behavioural changes (Beerda et al., 1999; Cozzi et al., 2016).

Shelters can represent a stressful environment for cats as well. Approximately 80% of Swedish shelters have experienced abnormal behaviours in sheltered cats, such as fearfulness, aggression, feeding disorders and inappropriate elimination behaviours (Eriksson et al., 2009). Moreover, as Gourkow et al. (2014) observed, sheltered cats display behavioural problems, such as crawling, freezing, feeling startled and retreating from humans – signs of poor welfare. These behaviours were found to have reduced their resistance to upper respiratory tract infections (Gourkow et al., 2013). Upper respiratory diseases represent the primary health issue reported in cats during their stay in shelters, supporting the hypothesis that behavioural elements and activities could be related to a poor health status (Gourkow et al., 2013).

The present work aims to evaluate the level of genomic damage in buccal mucosa cells of both shelter and family cats and dogs by buccal micronucleus assay. The tested hypothesis was that physiological stress conditions, like those potentially present in some shelters, could affect the levels of genomic damage in terms of increased frequencies of micronuclei (MNi), nuclear buds (NBUDs) and other nuclear rearrangements.

Buccal micronucleus assay is one of the most widely non-invasive techniques used to measure genetic damage in human and animal population studies (Lazalde-Ramos et al., 2017; Benvindo-Souza et al., 2019; Borges et al., 2019). MNi are chromosome fragments or whole chromosomes that fail to segregate properly during mitosis which appear in interphase as small additional nuclei. NBUDs are elimination processes from cells of amplified DNA and/or excess chromosomes (Fenech et al. 2011). It has been observed that the natural MNi frequency varies between certain limits (ranging from 3 to 23 MNi per 1000 cells) in different human populations. However, no frequency data is present in literature with regard to the prevalence of micronuclei in mammals like cats and dogs. In this scenario, the further purpose of our work was to evaluate, in buccal cells of these two mammals, the background level of genomic damage in terms of micronuclei and nuclear buds frequencies.

## 2. Materials and Methods

### 2.1. Subjects

The study population included thirty healthy mixed breed cats and thirty healthy mixed breed dogs, randomly sampled with a minimum two-year stay in a shelter, time that we consider sufficient for genomic damage to occur. As control groups, we selected healthy house cats (n = 30) and dogs (n = 30), using age/sex matching. Shelters were located in Turin, Piedmont, in Northwest Italy. All subjects were fed canned and/or packaged foods.

In order to evaluate the possible influence of the sex on the level of genomic damage, age and sex data was collected. It is well known that drugs and X-rays can alter the level of genomic damage (Santovito et al. 2017). Therefore, we excluded subjects that had contracted acute infections and/or chronic non-infectious diseases and exposure to diagnostic X-rays for a minimum of two years prior to the analysis.

All animals were treated and housed in compliance with Italian guidelines (available on http://www.aclonlus.org/wp-content/uploads/2014/02/LINEE-GUIDA-LR-34-97.pdf).

### 2.2. MNi assay

Exfoliated buccal mucosa cells were collected by gently scraping the mucosa of the inner lining of one or both cheeks with a spatula. Buccal cells were also collected from the inner side of the lower lip and palate. Indeed, the variability in MNi frequency between these areas was found to be minimal for control subjects (Holland et al., 2008). The tip of the spatula was immersed in a fixative solution consisting of methanol/Acetic Acid 3:1, stored at 4°C prior the analysis. Successively, cells were collected by centrifugation, the supernatant was discarded and the pellet was dissolved in a minimal amount of fixative which was seeded on the slides to detect MNi by conventional staining with 5% Giemsa (pH 6.8) prepared in Sörensen buffer.

Microscopic analysis was performed at 1000X magnification on a light microscope. MNi, NBUDs and other nuclear rearrangements were scored in 1,000 cells with well-preserved cytoplasm per subject according to the established criteria for MNi evaluation (Thomas et al., 2009).

### 2.3 Statistical Analysis

Statistical analyses were conducted using the SPSS software statistical package programme (version 24.0, Inc., Chicago, Illinois, USA). Differences between shelter and family cats and dogs as well as between sexes were evaluated by Kruskal-Wallis test. The correlation between age and the level of genomic damage was evaluated by regression analysis, whereas multivariate analysis was performed to identify sub-groups according to age and sex score. All *P*-values were two-tailed and the *a priori* level of statistical significance was set at P<0.05 for all tests.

## 3. Results

In Table 1 demographic characteristics of groups studied were reported. We sampled sixty cats, subdivided into thirty family cats (mean age 5.60±4.42, fourteen males and sixteen females) and thirty shelter cats (mean age 5.60±4.42, fifteen males and fifteen females). Similarly, for dogs, we sampled sixty subjects subdivided into thirty family dogs (mean age 6.40±3.73, twelve males and eighteen females) and thirty shelter dogs (mean age 5.41±1.64, eighteen males and twelve females). In both species, no significant differences were found between family and shelter subjects in terms of mean age.

**Table 1.** General characteristics of cat and dog samples.

|  | **Cats** | **Dogs** |
| --- | --- | --- |
| **Family** |  |  |
| N | 30 | 30 |
| Age (mean±SD) | 5.60±4.42 | 6.40±3.73 |
| Males | 14 | 12 |
| Females | 16 | 18 |
| **Shelter** |  |  |
| N | 30 | 30 |
| Age (mean±SD) | 5.23±4.43 | 5.41±1.64 |
| Males | 15 | 18 |
| Females | 15 | 12 |

N = number of studied subjects;
S.D. = Standard Deviation

In Table 2 results of the statistical evaluation of genomic damage between shelter and family cats and dogs were reported. In Figure 1 some examples of damaged cells observed in our samples were reported. Among family cats, the frequency of MNi, NBUDs and rearrangements were 0.100±0.383, 0.110±0.092, 0.008±0.119, with a frequency of total aberration of 0.287±0.405. Among shelter cats, the frequency of MNi, NBUDs and rearrangements were 0.210±0.209, 0.220±0.183, and 0.087±0.125, with a frequency of total aberration of 0.402±0.403. Significant differences were found between family and shelter cats in terms of MNi (*P*<0.001), NBUDs (*P* = 0.010) and total aberrations (*P* = 0.003).

**Table 2.** Statistical evaluation of genomic damage between Shelter and Family cats and dogs.

|  | N | Number of Analyzed Cells | MNi<br>N (Mean±SD %) | NBUDs<br>N (Mean±SD %) | REAR<br>N (Mean±SD %) | TOTAL ABERRATIONS<br>N (Mean±SD %) |
| --- | --- | --- | --- | --- | --- | --- |
| **CATS** |  |  |  |  |  |  |
| Family | 30 | 30,000 | 30 (0.100±0.383) | 33 (0.110±0.092) | 3 (0.008±0.119) | 86 (0.287±0.405) |
| Shelter | 30 | 30,000 | 63 (0.210±0.209)* | 66 (0.220±0.183)** | 26 (0.087±0.125) | 155 (0.402±0.403)*** |
| **DOGS** |  |  |  |  |  |  |
| Family | 30 | 30,000 | 25 (0.083±0.095) | 39 (0.130±0.154) | 12 (0.040±0.068) | 76 (0.253±0.229) |
| Shelter | 30 | 30,000 | 90 (0.300±0.234)* | 84 (0.280±0.220)* | 27 (0.090±0.092) | 201 (0.670±0.323)* |

N = number of studied subjects; S.D. = Standard Deviation; MNi = micronuclei; NBUDs = nuclear buds; REAR = rearrangements
\* $P < 0.001$ (Mann-Whitney and ANOVA tests) and $P = 0.029$ (Multivariate analysis) with respect to family group.
\*\* $P = 0.010$ (Mann-Whitney test).
\*\*\* $P = 0.003$ . with respect to family group (Mann- Whitney test), with respect to family group .

**Figure 1.**
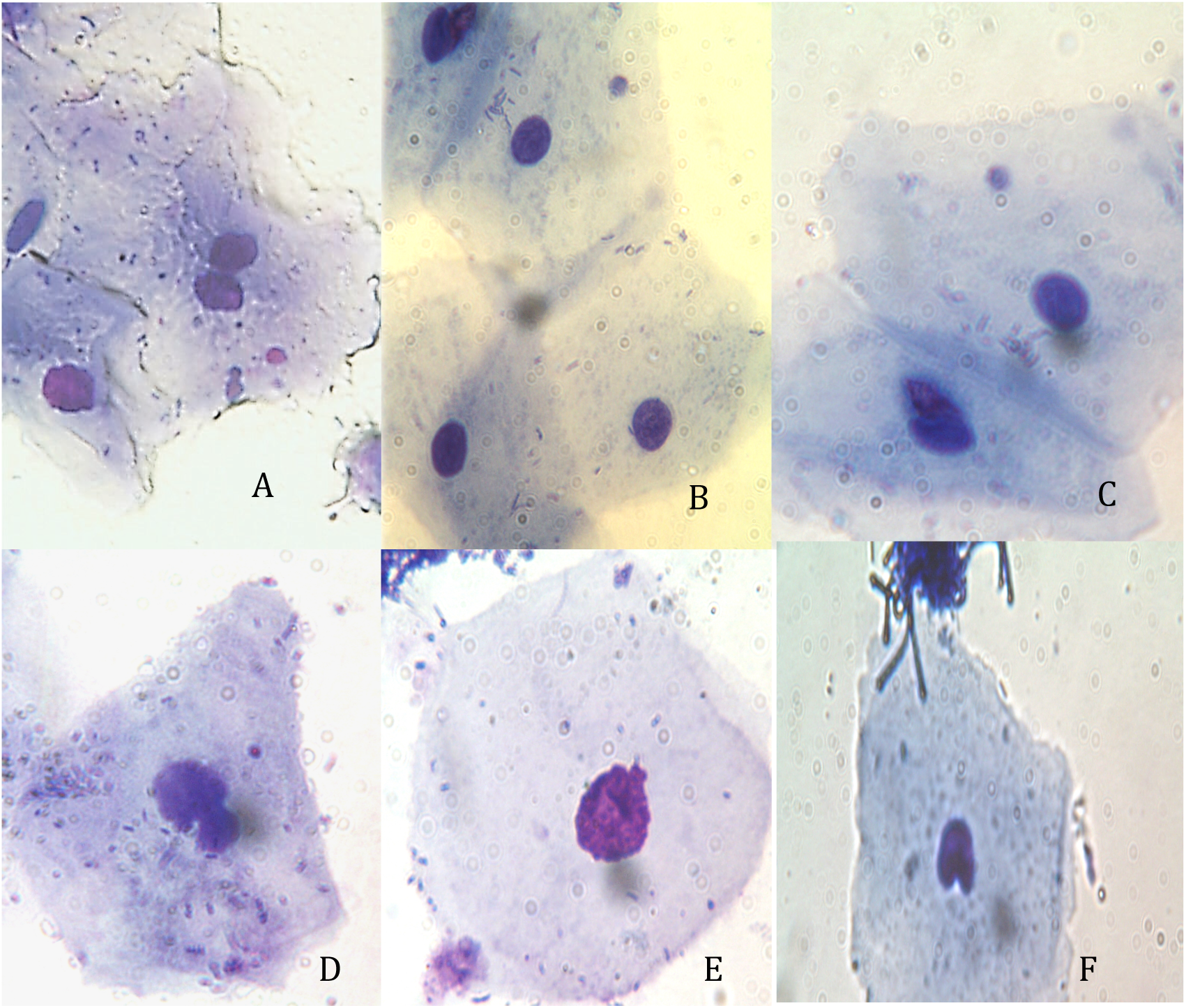
Examples of damaged cells observed in our samples. A) Binucleated cell with micronucleus; B) and C) mononucleated cells with micronucleus; D), E) nuclear buds; F) identation. These last two aberrations were included in the Rearrangement category.

Among dogs, the frequencies of MNi, NBUDs and rearrangements found in the family group were 0.083±0.095, 0.130±0.154, 0.040±0.068 with a frequency of total aberration of 0.253±0.229, whereas those observed among shelter dogs were 0.300±0.234, 0.280±0.220, 0.090±0.092 with a frequency of total aberration of 0.670±0.323. Significant differences were found between family and shelter dogs in terms of MNi, NBUDs and total aberrations (*P*<0.001).

In both species, no significant differences were found between sexes in terms of MNi, NBUds, rearrangement and total aberration frequencies (Table 3).

**Table 3.**
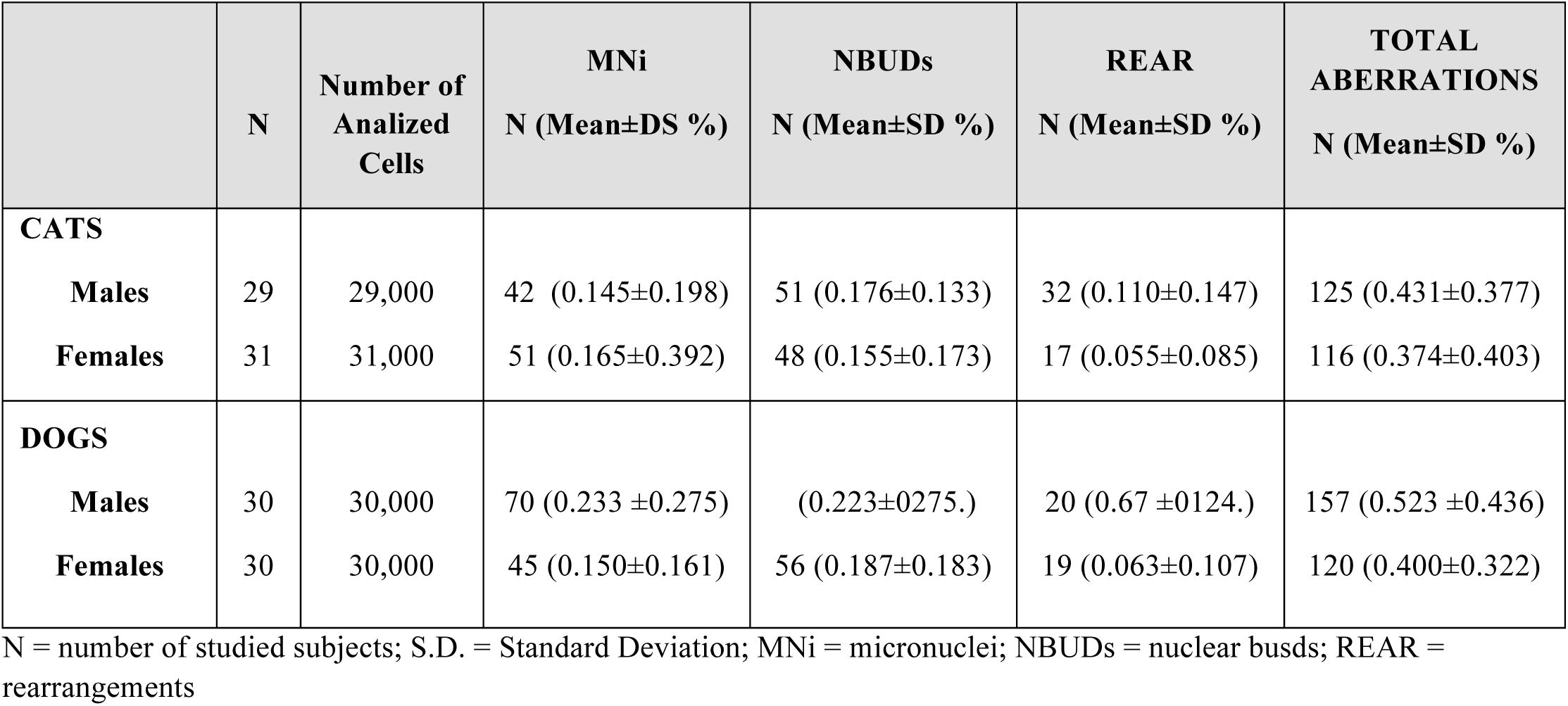
Evaluation of the level of genomic damage according to sex.

|  | N | Number of<br>Analyzed<br>Cells | MNi<br>N (Mean±DS %) | NBUDs<br>N (Mean±SD %) | REAR<br>N (Mean±SD %) | TOTAL<br>ABERRATIONS<br>N (Mean±SD %) |
| --- | --- | --- | --- | --- | --- | --- |
| **CATS** |  |  |  |  |  |  |
| **Males** | 29 | 29,000 | 42 (0.145±0.198) | 51 (0.176±0.133) | 32 (0.110±0.147) | 125 (0.431±0.377) |
| **Females** | 31 | 31,000 | 51 (0.165±0.392) | 48 (0.155±0.173) | 17 (0.055±0.085) | 116 (0.374±0.403) |
| **DOGS** |  |  |  |  |  |  |
| **Males** | 30 | 30,000 | 70 (0.233 ±0.275) | (0.223±0275.) | 20 (0.67 ±0124.) | 157 (0.523 ±0.436) |
| **Females** | 30 | 30,000 | 45 (0.150±0.161) | 56 (0.187±0.183) | 19 (0.063±0.107) | 120 (0.400±0.322) |

N = number of studied subjects; S.D. = Standard Deviation; MNi = micronuclei; NBUDs = nuclear busds; REAR = rearrangements

Finally, the regression analysis failed (P>0.05) to find a significant correlation between age and the frequencies of genomic markers. Similarly, the multivariate analysis did not show significantly any differences among the sub-groups according to age and sex (sex*age, *P* = 0.131 for cats and *P* = 0.988 for dogs)

## 4. Discussion

Domestic cats (*Felis silvestris catus*) and dogs (*Canis lupus familairis*) are two of the most popular companion animals in Western Countries. In Italy, in 2015, there were an estimated 1,051 authorized shelters housing more than 100,000 dogs and cats (Italian Health Ministry, 2015), whereas, in the U.S., approximately six to eight million cats and dogs enter shelters each year (HSUS, 2014). Shelters provide potentially aversive and stressful social environments, which in combination with the high turnover of animals contribute to the transmission of infectious diseases (Cohn, 2011; Hirsch, 2016)

To assess the possible influence of physiological stress on the level of genomic damage, we decided to evaluate the frequencies of MNi and other nuclear abnormalities in a sample of shelter cats and dogs and compare them with the levels of family cats and dogs.

Significant differences were found between shelter and family cats and dogs in terms of MNi, NBUDs and total rearrangements, which indicate that a condition of physiological stress, as can be observed in some shelters, may induce a high level of genomic damage.

The relationship between physiological stress and disease development was documented. There appears to be a significant connection between stress and immune responsiveness. When chronic, stress can weaken the immune system, causing disease susceptibility and the development of genomic damage (Gourkow *et al*., 2013). At genomic level, stress in mice and rats may induce alterations in the expression of hepatic gene, an up-regulation of several markers related to oxidative stress and an increase in apoptotic processes (Depke et al., 2009). Similarly, stress has been shown to influence brain DNA repair genes expression in rats whereas, stress, anxiety and depression have been shown to alter the methylation pattern of DNA in humans. Interestingly, it has been shown that stress caused by trauma increases genomic damage in humans. Children who have experienced violence have shown a significantly higher level of telomere erosion than their peers (Shalev et al., 2013; Bergholz et al., 2017; Kader et al., 2018).

Hence, a possible relationship between stressful conditions and increased frequencies of MNi is not surprising. In humans, higher levels of MNi in peripheral blood lymphocytes and other cell types have been associated, in perspective, with an increased risk of cancer (Bonassi et al., 2011). Similarly, we cannot rule out a connection between higher levels of MNi and a higher incidence of cancer even in cats and dogs living in shelters as compared to family cats and dogs.

In addition, MNi do not represent only the products of biological errors, but trigger the activation of the immune system related genes through the exposure of DNA fragments, which suggests that the presence of MNi can be perceived by the immune system (Gekara, 2017). MNi also represent a mechanism of elimination of genetic material, such as amplified genes, and contribute to nuclear dynamics and genomic chaos (Heng 2019; Ye et al., 2019). The latter represents a process of rapid genomic re-organization that results in the formation of very altered and chaotic genomes (defined by both extreme structural and numerical alterations), some of which can be selected to establish stable genomes (Ye et al., 2019).

Finally, in contrast to Santovito et al. (2020), we found no effect of age on the level of genomic damage neither on dogs nor on cats. It is plausible that the relatively short life expectancy of these two species may mask any possible correlation between age and MNi frequency.

## 5. Conclusions

In this work we provided evidence of a possible correlation between physiological stress conditions and higher levels of genomic damage in a sample of sheltered cats and dogs. However, we wish to underline that the results of this study cannot be generalized to all animal shelters as we are aware that some animal shelters offer a comfortable place, in terms of space and care. Our work will hopefully serve as a stimulus for those shelters that, for various reasons, are unable to provide a relaxing environment for animals. Furthermore, given the relatively low cost of laboratory procedures, these techniques, combined with more traditional ones, such as behavioural tests, could provide a more comprehensive picture of the health status of animal communities.

## Acknowledgements

The authors would like to thank all veterinary and shelter volunteers who allowed us access to shelters and participated in the collection of buccal samples and that offer valuable work useful to improve animal welfare. We would also like to thank the Professor Sonia Slaviero for her contribution in revising English.

This study was financed by University of Turin with local 2015-2018 grants.

## References

1. Beerda, B., Schilder, M.B.H., Bernardina, W., Van Hoof, J.A.R.A.M., De Vries H.W., Mol, J.A. (1999). Chronic stress indogs subjected to social and spatial restriction. I: Behavioral responses. Physiological Behavior, 66, 233–242.

2. Beerda, B., Schilder, M.B.H., Van Hoof, J.A.R.A.M., De Vries, H.W., Mol, J.A. (2000). Behavioral and hormonal indicators of enduring environmental stress in dogs. Animal Welfare, 9, 49–62.

3. Benvindo-Souza, M., Borges, R.E., Pacheco, S.M., de Souza Santos, L.R. (2019). Genotoxicological analyses of insectivorous bats (Mammalia: Chiroptera) in central Brazil: The oral epithelium as an indicator of environmental quality. Environmental Pollution, 245, 504–509.

4. Bergholz, L.M., Subic-Wrana, C., Heylmann, D., Beutel, M.E., Wiltink, J., Kaina, B. (2017). DNA damage in lymphcytes of patients suffering from complex traumatization. DNA Repair, 52, 103–109.

5. Bonassi, S., El-Zein, R., Bolognesi, C, Fenech, M. (2011). Micronuclei frequency in peripheral blood lymphocytes and cancer risk: evidence from human studies. Mutagenesis, 26, 93–100.

6. Borges, R.E., de Souza Santos, L.R., Benvindo-Souza, M., Modesto, R.S., Assis, R.A., de Oliveira, C. (2019). Genotoxic Evaluation in Tadpoles Associated with Agriculture in the Central Cerrado, Brazil. Archives of Environmental Contamination Toxicology, 77(1), 22–28.

7. Cohn, L.A. (2011). Feline Respiratory Disease Complex. Veterinary Clinics of North America –Small Animal Practice, 41(6), 1273–1289.

8. Coppola, C.L., Grandin, T., Enns, R.M. (2006). Human interaction and cortisol: Can human contact reduce stress for shelter dogs? Physiological Behavior, 87: 537–541.

9. Cozzi, A., Mariti, C., Ogi, A., Sighieri, C., Gazzano, A. (2016). Behavioral modification in sheltered dogs. Dog Behaviour, 3, 1–12.

10. Depke, M., Steil, L., Domanska, G., Völker, U., Kiank, G. (2009). Altered hepatic mRNA expression of immune response and apoptosis-associated genes after acute and chronic psychological stress in mice. Molecular Immunology, 46(15), 3018–3028.

11. Eriksson, P., Loberg, J., Andersson, M. (2009). A survey of cat shelters in Sweden. Animal Welfare, 18(3), 283–288.

12. Fenech, M., Kirsch-Volders, M., Natarajan, A.T., Surralles, J., Crott, J.W., Parry, J., Norppa, H., Eastmond, D.A., Tucker, J.D., Thomas, P. (2011). Molecular mechanisms of micronucleus, nucleoplasmic bridge and nuclear bud formation in mammalian and human cells. Mutagenesis, 26, 125–132.

13. Gekara, N.O. (2017). DNA damage-induced immune response: Micronuclei provide key platform. The Journal of Cell Biology, 216, 2999–3001.

14. Gourkow, N., Lawson, J.H., Hamon, S.C., Phillips, C.J.C. (2013). Descriptive epidemiology of upper respiratory disease and associated risk factors in cats in an animal shelter in coastal western Canada. The Canadian Veterinary Journal, 54(2), 132–138.

15. Gourkow, N., LaVoy, A., Dean, G., Phillipes, C. (2014). Associations of behaviour with secretory immunoglobulin A and cortisol in domestic cats during their first week in an animal shelter. Applied Animal Behaviour Science, 150, 55–64.

16. Heng, H.H. (2019). Genome Chaos: Rethinking Genetics, Evolution, and Molecular Medicine; Academic Press Elsevier: Cambridge, MA, USA, 2019; ISBN 978-012-8136-35-5.

17. Hennessy, M.B., Voith, V.L., Mazzei, S.J., Buttram, J., Miller, D.D., Linden, F. (2001). Behavior and cortisol levels of dogs in a public animal shelter, and an exploration of the ability of these measures to predict problem behavior after adoption. Applied Animal Behaviour Science, 73: 217–233.

18. Hirsch, E.N. (2016). Feline Stress. Methodological considerations for non-invasive assessment of cats housed in groups and singly. PhD thesis, Swedish University of Agricultural Sciences, Skara. Available on https://pub.epsilon.slu.se/13682/1/hirsch_en_160927.pdf. Accessed on 10-29-20191.

19. Holland, N., Bolognesi, C., Kirsch-Volders, M., Bonassi, S., Zeiger, E., Knasmueller, S., Fenech, M. (2008). The micronucleus assay in human buccal cells as a tool for biomonitoring DNA damage: the HUMN project perspective on current status and knowledge gaps. Mutation Research, 659(1-2), 93–108.

20. Humane Society of the United States (HSUS) (2014). Annual Report 2014. Available on: https://www.humanesociety.org/sites/default/files/docs/2014-hsus-annual-report.pdf. Accessed on 11-10-2019.

21. Italian Health Minister (2015). Cani e Rifugi. Available on: http://www.salute.gov.it/portale/temi/p2_6.jsp?lingua=italiano&id=3093&area=cani&menu=abbandono). Accessed on 11-10-2019.

22. Kader, F., Ghai, M., Maharaj, L. (2018). The effects of DNA methylation on human psycology. Behavioural Brain Research, 346, 47–65.

23. Kessler, M.R. & Turner, D.C. (1999). Effects of density and cage size on stress in domestic cats (Felis silvestris catus) housed in animal shelters and boarding catteries. Animal Welfare, 8(3), 259–267.

24. Lazalde-Ramos, B.P., Zamora-Péres, A.L., Sosa-Macía, M., Galaviz-Hernández, C., Zúñiga-González, G.M. (2017). Micronculei and nuclear anomalies in Mexico’s indigenous population. Salud Publica Mexico, 59, 532–539.

25. Righi, C., Menchetti, L., Orlandi, R., Moscati, L., Mancini, S., Diviero, S. (2019). Welfare Assessment in Shelter Dogs by Using Physiological and Immunological Parameters. Animals (Basel), 9(6), 340.

26. Santovito, A., Delsoglio, M., Manitta, E., Picco, G., Meschiati, G., Chiarizio, M., Gendusa, C., Cervella, P. (2017). Association of GSTT1 Null, XPD 751 CC and XPC 939 CC Genotypes With Increased Levels of Genomic Damage Among Hospital Pathologists. Biomarkers, 22(6), 557–565.

27. Santovito, A., Gendusa, C. (2020). Micronuclei Frequency in Peripheral Blood Lymphocytes of Healthy Subjects Living in Turin (North-Italy): Contribution of Body Mass Index, Age and Sex. Annals of Human Biology, 47(1), 48–54.

28. Shalev, I., Moffitt, T.E., Sugden, K., Williams, B., Houst, R.M., Danese, A., Mill, J., Arseneault L., Caspi, A. (2013). Exposure to violence during childhood is associated with telomere erosion from 5 to 10 years of age: a longitudinal study. Molecular Psychiatric, 18(5), 576–581.

29. Taylor, K.D., Mills, D.S. (2007). The effect of the kennel environment on canine welfare: A critical review of experimental studies. Animal Welfare, 16, 435–447.

30. Thomas, P., Holland, N., Bolognesi, C., Kirsch-Volders, M., Bonassi, S., Zeiger, E., Knasmueller, S., Fenech, M. (2009). Buccal micronucleus cytome assay. Nature Protocols, 4(6), 825–837.

31. Titulaer, M., Blackwell, E.J., Mendl, M., Casey, R.A. (2013). Cross sectional study comparing behavioural, cognitive and physiological indicators of welfare between short and long term kennelled domestic dogs. Applied Animal Behavior Science, 147, 149–158.

32. Ye, C.J., Sharpe, Z., Alemata, S., Mackenzie, S., Liu, G., Abdallah, B., Horne, S., Regan, S., Heng, H.H. (2019). Micronuclei and Genome Chaos: Changing the System Inheritance. Genes (Basel), 10(5):366.

33. Wells, D.L., Hepper, P.G. (2000) Prevalence of behavior problems reported by owners of dogs purchased from animal rescue. Applied Animal Behavior Science, 69, 55–65.

34. Wells, D.L., Graham, L., Hepper, P.G. (2002). The influence of auditory stimulation on the behavior of dogs housed in a rescue shelter. Animal Welfare, 11: 385–393.

